# Molecular basis for the pheromone-mediated feeding preference of early-instar *Mythimna separata* larvae

**DOI:** 10.1101/2022.06.29.498063

**Authors:** Jiyuan Liu, Shichang Cheng, Tong Zhou, Ruichi Li, Zhen Tian

## Abstract

Insect sex pheromones are important chemical signals in sexual communication, they are traditionally thought to be inactive to larvae. However, it was found that some lepidopteran larvae can be significantly attracted by sex pheromones of the same species in a food context. Here we reported that the host-feeding choice of early-instar *Mythimna separata* larvae can be effectively regulated by the sex pheromone component (*Z*)-11-hexadecenal (*Z*11-16: Ald). Further exploration revealed that two olfactory proteins, *M. separata* general odorant binding protein 2 (MsGOBP2) and olfactory receptor 3 (MsOR3), were molecular basis of the host-feeding choice regulation mediated by *Z*11-16: Ald. Silencing either *MsGOBP2* or *MsOR3* led to a sharply reduced preference for *Z*11-16: Ald-spiked hosts. It is possible that the *Z*11-16: Ald-based preference of the *M. separata* larvae for host is governed by the interplay between MsGOBP2 and MsOR3. In the present research, the interactions between MsGOBP2 and *Z*11-16: Ald were also discussed using molecular dynamics-based approaches. Our research explored insight into the *Z*11-16: Ald-mediated host-feeding choice regulation of *M. separata* larvae, all the results would aid in developing olfaction-based methods for controlling pests in larval stage.

## 1 INTRODUCTION

Insects use olfactory cues to sense the outside world, many of their fundamental behaviors including feeding, mating, and oviposition are regulated by chemical signals^[1, 2]^. Insects have developed sophisticated olfactory systems of extraordinary specificity and sensitivity to detect these chemical stimuli^[3-5]^. In the olfactory events of insects, at least two classes of proteins are required, olfaction receptors (ORs) and odorant binding proteins (OBPs)^[6-9]^. As a family of small soluble proteins, OBPs are known to bind semiochemicals (pheromones and odorants) and ferry them to ORs expressed on the dendrites of specific olfactory sensory neurons (OSNs)^[1, 10, 11]^. Tremendous evidences have revealed the enhanced, sometimes more specific, responses of ORs to semiochemicals in the presence of OBPs^[12-15]^. Specific OBPs recognizing particular semiochemicals have also been identified in more than one insect species^[16-19]^. The indispensability of OBPs in insect olfaction made it an attractive target for developing behaviorally active semiochemcials. These days, reverse chemical ecology, an efficient method of discovering semiochemcials based on their binding affinity to olfactory proteins including OBPs, has been widely used in identifying attractants and repellents for more than one insect species^[20-23]^.

Female-released sex pheromones are important chemical signals in intraspecific sexual communication, they are usually multicomponent mixtures with specific ratios, contributing to the accurate recognition of females by males of the same species^[24, 25]^. As one of the best-studied systems, an acceptable mechanism for pheromone signaling have been demonstrated in quite a few lepidopteran insects (moths and butterflies). With the help of pheromone binding proteins (PBPs), a subclass of OBPs, sex pheromone molecules were ferried to the membrane-bound pheromone receptors (PRs). Then the transduced sex pheromone signals were transmitted to the macroglomerular complex in antennal lobes, integrated in brain centers, and finally translated into behavioral responses^[25-27]^. It is clear that PBPs, as their name implies, are necessary to the activation of sex pheromone signals transduction in lepidopteran adults^[1, 3]^.

Recent studies showed that the food selection of some lepidopteran larvae can be affected by the species-specific sex pheromones in the food context. Take the larvae of *Spodoptera litura, S. littoralis* and *Plutella xylostella* for example, they were more attracted by the food containing their sex pheromones^[28-30]^. On this basis, the species-specific sex pheromones were regarded as an olfactory cue guiding larvae to the better food. This is contrary to the traditional view that lepidopteran larvae are not necessarily able to detect sex pheromones due to sexual immaturity. Since no genes encoding PBPs are expressed at the larval stage, the sense of sex pheromone in larvae is believed to be mediated by general odorant binding proteins (GOBPs)^[28, 30]^, a second subclass of OBPs abundantly expressed in larval antennae. More than one researches have detailed the binding of lepidopteran GOBPs to their species-specific sex pheromones^[28, 31-35]^.

The oriental armyworm, *Mythimna separata* (Walker) is an important lepidopteran pest causing serious damages to cereal crops (rice, wheat, maize, etc.) in North China^[36, 37]^. Its major sex pheromone component, (*Z*)-11-hexadecenal (*Z*11-16: Ald), has been developed as an efficient tool for controlling and monitoring the pest^[26, 38]^. The perception mechanism of *Z*11-16: Ald at adult level has been clearly revealed as well, where the *M. separata* OR1 (MsOR1) and *M. separata* OR3 (MsOR3) were verified as two key proteins bridging the intracellular and extracellular signal transduction^[25, 26, 39]^. In comparison, the chemoreception system of *M. separata* larvae has received little attention, although larval stage is the most damaging.

In this study, we reported an enhanced response to hosts mediated by *Z*11-16: Ald in the early-instar *M. separata* larvae. To get insight into the *Z*11-16: Ald-based preference for host leaves, molecular basis of *Z*11-16: Ald perception by *M. separata* larvae were explored as well. Then, molecular dynamics-based approaches were employed to uncover interactions between MsGOBP2 and *Z*11-16: Ald. Identifying the role of sex pheromone in attracting early-instar *M. separata* larvae and revealing the larval perception mechanism of sex pheromone would provide basis for olfaction-based management of *M. separata* larvae.

## 2 RESULTS

A two-choice test was used to test the effects of *Z*11-16: Ald, the major component of *M. separata* sex pheromone, on the host-feeding choice of *M. separata* larvae. It can be seen that *Z*11-16: Ald alone showed no attractivity to the 1st-4th instar *M. separata* larvae (Figure S1). Further tests revealed that fresh maize leaves spiked with *Z*11-16: Ald were preferred by the 1st (χ^2^=28.80, *P*<0.01) and the 2nd (χ^2^=70.56, *P*<0.01) instar larvae. As shown in Figure 1A, most early-instar larvae (1st-2nd instar) chose the “maize leaf + *Z*11-16: Ald” food source. While for the 3rd instar larvae (χ^2^=1.09, *P*>0.05), their feeding preference for fresh maize leaves was not evidently regulated by *Z*11-16: Ald (Figure 1A).

**Figure 1.**
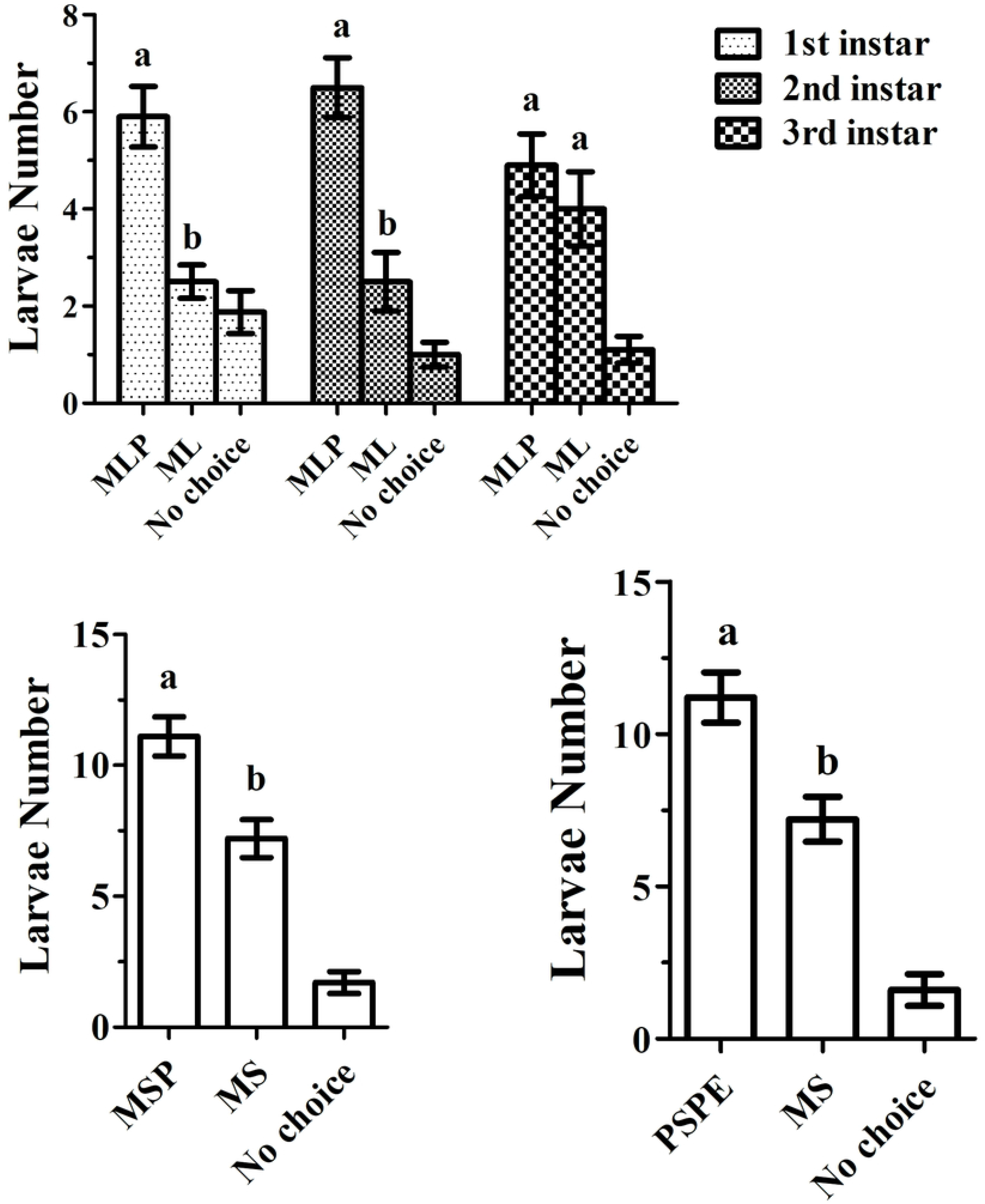
Influences of *Z*11-16: Ald on the host-feeding choice of the *Mythimna separata* larvae. MLP means maize leaf (diameter=1.5 cm) spiked with the sex pheromone component *Z*11-16: Ald, ML means maize leaf spiked with the GC-grade solvent methanol, MSP means maize seedling spiked with *Z*11-16: Ald, MS means maize seedling spiked with methanol, PSPE means pea seedling spiked with *Z*11-16: Ald and crude extracts of maize leaves. No choice means *M. separata* larvae make no choice within test time. Columns marked with different letters in a group mean significant difference.

We further tested the host-feeding choice regulation activity of *Z*11-16: Ald using maize seedlings. It is clear that the 2nd instar larvae can be significantly attracted by the maize seedlings sprayed with *Z*11-16: Ald (Figure 1B). The pea seedlings (non-host of *M. separata* larvae) sprayed with crude extracts of fresh maize leaves became attractive to the 2nd instar *M. separata* larvae (Figure S2), with the larvae-attracting capacity being similar to that of maize seedlings. However, the pea seedlings sprayed with crude extracts can be mistaken for a better host by the 2nd instar *M. separata* larvae when appropriate *Z*11-16: Ald was present. As shown in Figure 1C, the number of *M. separata* larvae attracted by treated pea seedlings (sprayed with maize leaf crude extracts and *Z*11-16: Ald) was significantly higher than that attracted by maize seedlings. The influence of *Z*11-16: Ald on the host-feeding choice of early-instar larvae suggested that the sex pheromone *Z*11-16: Ald can be perceived by *M. separata* larvae.

Now that *Z*11-16: Ald are active to enhance the responses of early-instar *M. separata* larvae to hosts, it would be interesting to make clear the molecular basis of the *Z*11-16: Ald-mediated food preference. We initially determined the expression profiles of 10 olfactory genes reported in *M. separata* adults, including three *MsPBPs* (*MsPBP, MsPBP1* and *MsPBP2*), two *MsGOBPs* (*MsGOBP1* and *MsGOBP2*), two pheromone receptors (*MsPR1* and *MsPR2*), and three odorant receptors (*MsOR1, MsOR2* and *MsOR3*). The results of RT-PCR plus gene sequencing showed that no *MsPBPs* and *MsOR1* genes were detected in the larvae, but genes encoding MsGOBP1, MsGOBP2, MsPR1, MsOR2 and MsOR3 proteins were expressed (Figure 2). Although weak amplification bands can be observed in the *MsPBP* and *MsPBP1* lanes (Figure 2), further cloning and sequencing verified them as unrelated genes.

**Figure 2.**
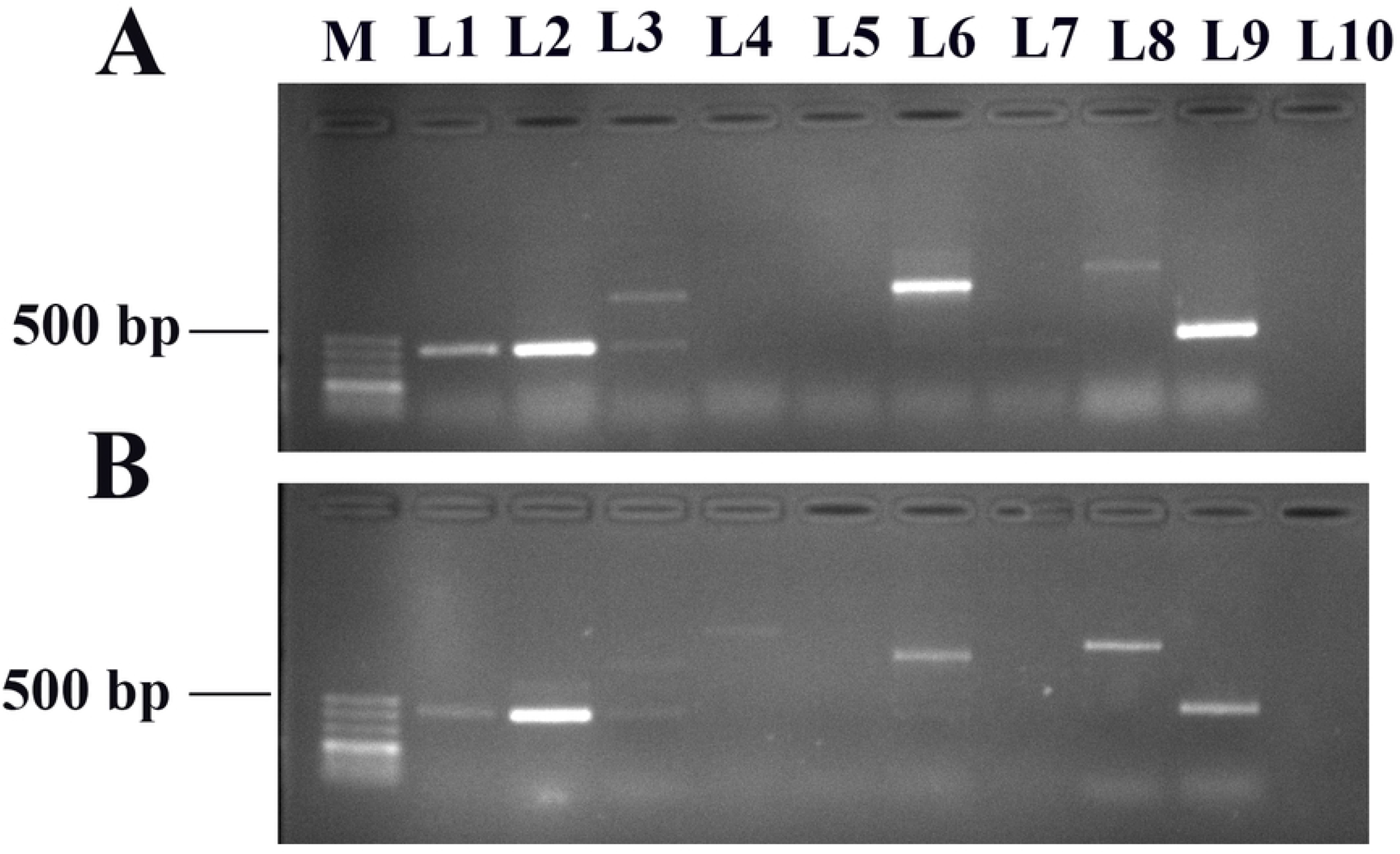
Expression profiles of ten olfactory genes in the 1st (A) and 2nd (B) instar *Mythimna separata* larvae. M is the 500 bp DNA ladder. L1-L10 are partial genes encoding MsGOBP1, MsGOBP2, MsPBP, MsPBP1, MsPBP2, MsPR1, MsPR2, MsOR2, MsOR3, and MsOR1. Weak amplification bands in the L3 and L4 lanes (MsPBP and MsPBP1) are verified as unrelated genes by sequencing.

We continued to explore the potential role of the two soluble proteins (MsGOBP1 and MsGOBP2) in the *Z*11-16: Ald recognition by early-instar *M. separata* larvae. The recombinant MsGOBP1 and MsGOBP2 proteins were expressed in the prokaryotic expression system (Figure S3) and used for *in vitro* binding assay. It is shown that, although both MsGOBPs can bind the molecule *Z*11-16: Ald, MsGOBP2 was provided with a relatively higher affinity, with *K*_d_ values being 1.94±0.37 μM for MsGOBP2 and 10.03±0.93 μM for MsGOBP1 (Figure 3).

**Figure 3.**
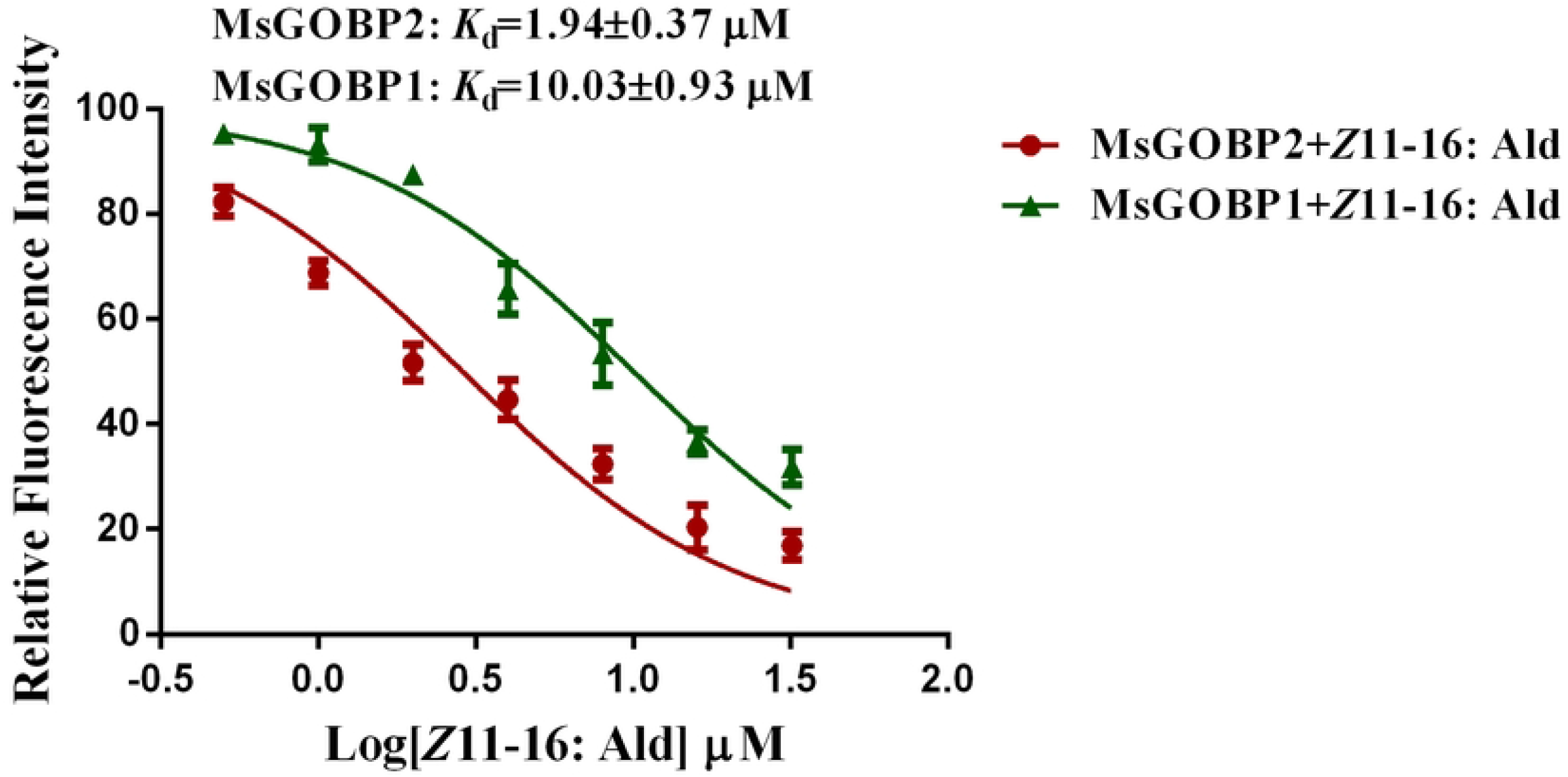
Dose-response curves for *Z*11-16: Ald against the two recombinant MsGOBPs.

RNAi was employed to silence the *in vivo* expression of the *MsGOBP1* and *MsGOBP2* genes. As shown in Figure 4A, expression levels of the two soluble olfactory proteins were evidently decreased by at least 50%. The preference of early-instar *M. separata* larvae for the host spiked with *Z*11-16: Ald was reduced to a varying degree when the expression of either MsGOBP1 or MsGOBP2 was suppressed. Especially for MsGOBP2, its decreased expression almost led to the disappearance of *Z*11-16: Ald-based host preference (Figure 4B). The decreased larval response to *Z*11-16: Ald caused by silencing MsGOBP2 showed no mitigation when increasing to 50 ng (data not show). Thus, in larval antennae, it is possible that MsGOBP2 plays a leading role and MsGOBP1 plays an auxiliary or invalid part in mediating the *Z*11-16: Ald-induced host preference.

**Figure 4.**
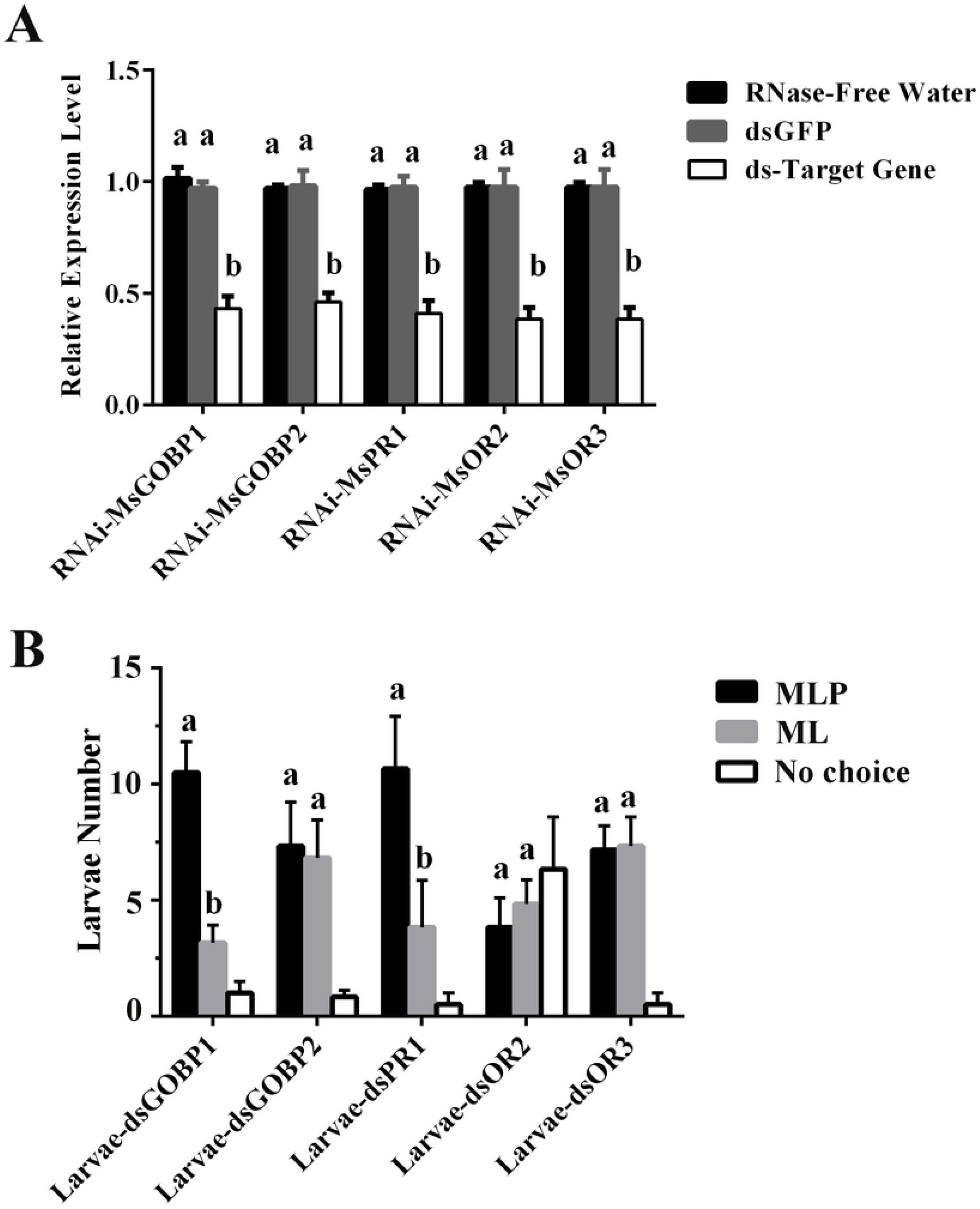
The silence efficiency of five target genes (A) and the effects of silencing individual target genes on the attraction of *Mythimna separata* larvae to maize leaves spiked with *Z*11-16: Ald (B). MLP means maize leaf (diameter=1.5 cm) spiked with the sex pheromone component *Z*11-16: Ald, ML means maize leaf spiked with the GC-grade solvent methanol, No choice means *M. separata* larvae make no choice within test time. Columns marked with different letters in a group mean significant difference. RNAi-MsGOBP1, RNAi-MsGOBP2, RNAi-MsPR1, RNAi-MsOR2 and RNAi-MsOR3 mean RNAi of genes encoding MsGOBP1, MsGOBP2, MsPR1, MsOR2 and MsOR3, respectively. Larvae-dsGOBP1, Larvae-dsGOBP2, Larvae-dsPR1, Larvae-dsOR2 and Larvae-dsOR3 mean *M. separata* larvae with genes encoding MsGOBP1, MsGOBP2, MsPR1, MsOR2 and MsOR3 being silenced, respectively.

We also emphatically checked whether the three olfaction receptors (MsPR1, MsOR2 and MsOR3) expressed in larval antennae were involved in the *Z*11-16: Ald-based preference for hosts. After continuously feeding the newly-hatched larvae with corresponding dsRNA for 72 h, the expression levels of *MsPR1, MsOR2* and *MsOR3* were decreased by around 60% (Figure 4A). Behavioral tests revealed that the *MsPR1*-silenced *M. separata* larvae kept a strong response to maize leaves spiked with *Z*11-16: Ald (Figure 4B). For *MsOR2*, its silence decreased the number of larvae attracted by both types of food (maize leaves with and without *Z*11-16: Ald) (Figure 4B). Considering the fact that *Z*11-16: Ald is attractive to early-instar *M. separata* larvae only in a food context, it’s unable to judge whether MsOR2 is involved in the larval perception of *Z*11-16: Ald. Larvae expressing a decreased level of *MsOR3* showed a greatly weakened response to *Z*11-16: Ald in a food context (Figure 4B). Combined RNAi and behavioral tests revealed the necessity of MsOR3 to the *Z*11-16: Ald-mediated host-feeding choice regulation in *M. separata* larvae.

Now that the soluble olfactory protein MsGOBP2 was key to the *in vivo* perception of *Z*11-16: Ald, we further discussed the MsGOBP2-*Z*11-16: Ald interactions at atomic level. The MsGOBP2-*Z*11-16: Ald complex was constructed by docking *Z*11-16: Ald into the cavity of MsGOBP2, and subsequently analyzed by MD-based approaches. MD simulations revealed that both the MsGOBP2-*Z*11-16: Ald system and the inclusive molecule *Z*11-16: Ald achieved equilibrium in quite a short time (Figure S8A&B). In the representative conformation produced by MD simulations (Figure 5A), it can be seen that the *Z*11-16: Ald molecule was stabilized in a hydrophobic pocket consisted of Thr9, Phe12, Phe36, Ser56, Ile52, Ile94, Val111 and Phe118. As principal residues interacting with the ligand *Z*11-16: Ald, free energy contributions of all these eight residues were fairly close to (≥ 0.95 kcal/mol) or above −1.00 kcal/mol (Figure 5B, Table 1). Especially for Phe12, Ile52 and Ile94, their total free energy contributions (T_TOT_) even exceeded −1.50 kcal/mol (Figure 4B, Table 1). Meanwhile, regions around these residues all exhibited fairly low flexibility during the MD process (Figure S8C). The results of per-residual free energy decomposition revealed that these eight residues all contributed vast amounts of van der Waals energy (ΔE_VDW_≥-0.70 kcal/mol). Correspondingly, ΔE_VDW_ (ΔE_VDW_=−46.78 kcal/mol) was the largest *in silico* binding free energy (ΔG_bind-cal_) contributor of the MsGOBP2-*Z*11-16: Ald complex (Table 2). Comparing with ΔE_VDW_, the ΔG_bind-cal_ item (Table 2) provided by electrostatic energy was fairly weak (ΔE_ELE_=-2.86 ± 0.24 kcal/mol). Among the residues dominantly interacting with *Z*11-16: Ald, Ser56, the residue forming a hydrogen-bond (3.0 Å) with the O of *Z*11-16: Ald carbonyl (Figure 4A), was the only residue contributing relatively higher ΔE_ELE_ (−0.91 kcal/mol) (Table 1). The cross-validation of *in vitro* binding assay, MD, binding free energy calculation, per-residual free energy decomposition indicated the high affinity between MsGOBP2 and *Z*11-16: Ald.

**Table 1.** The results of per-residue free energy decomposition

| Residue | S<sub>VDW</sub> | B<sub>VDW</sub> | T<sub>VDW</sub> | S<sub>ELE</sub> | B<sub>ELE</sub> | T<sub>ELE</sub> | S<sub>EPB</sub> | B<sub>EPB</sub> | T<sub>EPB</sub> | S<sub>TOT</sub> | B<sub>TOT</sub> | T<sub>TOT</sub> |
| --- | --- | --- | --- | --- | --- | --- | --- | --- | --- | --- | --- | --- |
| Thr9 | -0.51 | -0.39 | -0.90 | -0.15 | 0.02 | -0.13 | -0.02 | -0.12 | -0.14 | -0.69 | -0.52 | -1.21 |
| Phe12 | -1.74 | -0.14 | -1.88 | -0.007 | 0.027 | 0.02 | 0.28 | -0.45 | -0.17 | -1.65 | -0.28 | -1.93 |
| Phe36 | -0.96 | -0.04 | -1.00 | -0.02 | 0.00 | -0.02 | 0.18 | -0.07 | 0.11 | -0.85 | -0.11 | -0.96 |
| Ile52 | -1.14 | -0.29 | -1.43 | 0.01 | -0.06 | -0.05 | -0.07 | 0.15 | 0.08 | -1.35 | -0.2 | -1.55 |
| Ser56 | -0.54 | -0.16 | -0.70 | -0.80 | -0.11 | -0.91 | 0.76 | -0.01 | 0.75 | -0.67 | -0.28 | -0.95 |
| Ile94 | -1.33 | -0.13 | -1.46 | 0.08 | -0.03 | 0.05 | -0.12 | 0.03 | -0.09 | -1.56 | -0.13 | -1.69 |
| Val111 | -0.63 | -0.26 | -0.89 | -0.01 | -0.14 | -0.15 | -0.01 | 0.12 | 0.11 | -0.75 | -0.28 | -1.03 |
| Phe118 | -1.11 | -0.10 | -1.21 | -0.04 | 0.03 | -0.01 | 0.22 | -0.03 | 0.19 | -1.05 | -0.09 | -1.14 |

Note: Energies are shown as contributions from van der Waals energy (VDW), electrostatic energy (ELE), electrostatic contribution to the solvation free energy (EPB) and total energy (TOT) of sidechain atoms (S), backbone atoms (B), and the sum of them (T). All values are shown in kcal/mol.

**Table 2.** Theoretical and experimental binding free energy^*a*^ for the MsGOBP2-*Z*11-16: Ald complex

| System | $\Delta E_{ELE}^b$ | $\Delta E_{VDW}^c$ | $\Delta E_{EPB}^d$ | $\Delta E_{ECAVITY}^e$ | $\Delta G_{GAS}^f$ | $\Delta G_{SOL}^g$ | $\Delta G_{bind-cal}^h$ | $\Delta G_{bind-exp}^i$ |
| --- | --- | --- | --- | --- | --- | --- | --- | --- |
| MsGOBP2- | -2.86 | -46.78 | 22.78 | -34.15 | -49.64 | 43.71 | -5.93 | -7.66 |
| Z11-16: Ald | (0.24) | (0.20) | (0.26) | (0.07) | (0.31) | (0.28) | (0.38) |  |

Note: <sup>a</sup> All values are given in kcal/mol, with corresponding standard errors of mean in parentheses. <sup>b-g</sup> $\Delta E_{ELE}$ , $\Delta E_{VDW}$ , $\Delta E_{EPB}$ , $\Delta E_{ECAVITY}$ , $\Delta G_{GAS}$ , and $\Delta G_{SOL}$ are electrostatic energy, van der Waals energy, electrostatic contribution to the solvation free energy, nonpolar contribution to the solvation free energy, total gas phase free energy, and sum of non-polar and polar contributions to solvation,
respectively.
<sup>h</sup> The theoretical binding free energy calculated using the MM-PBSA method in Amber12.
<sup>i</sup> The experimental binding free energy calculated based on the dissociation constant between MsGOBP2 and Z11-16: Ald.

**Figure 5.**
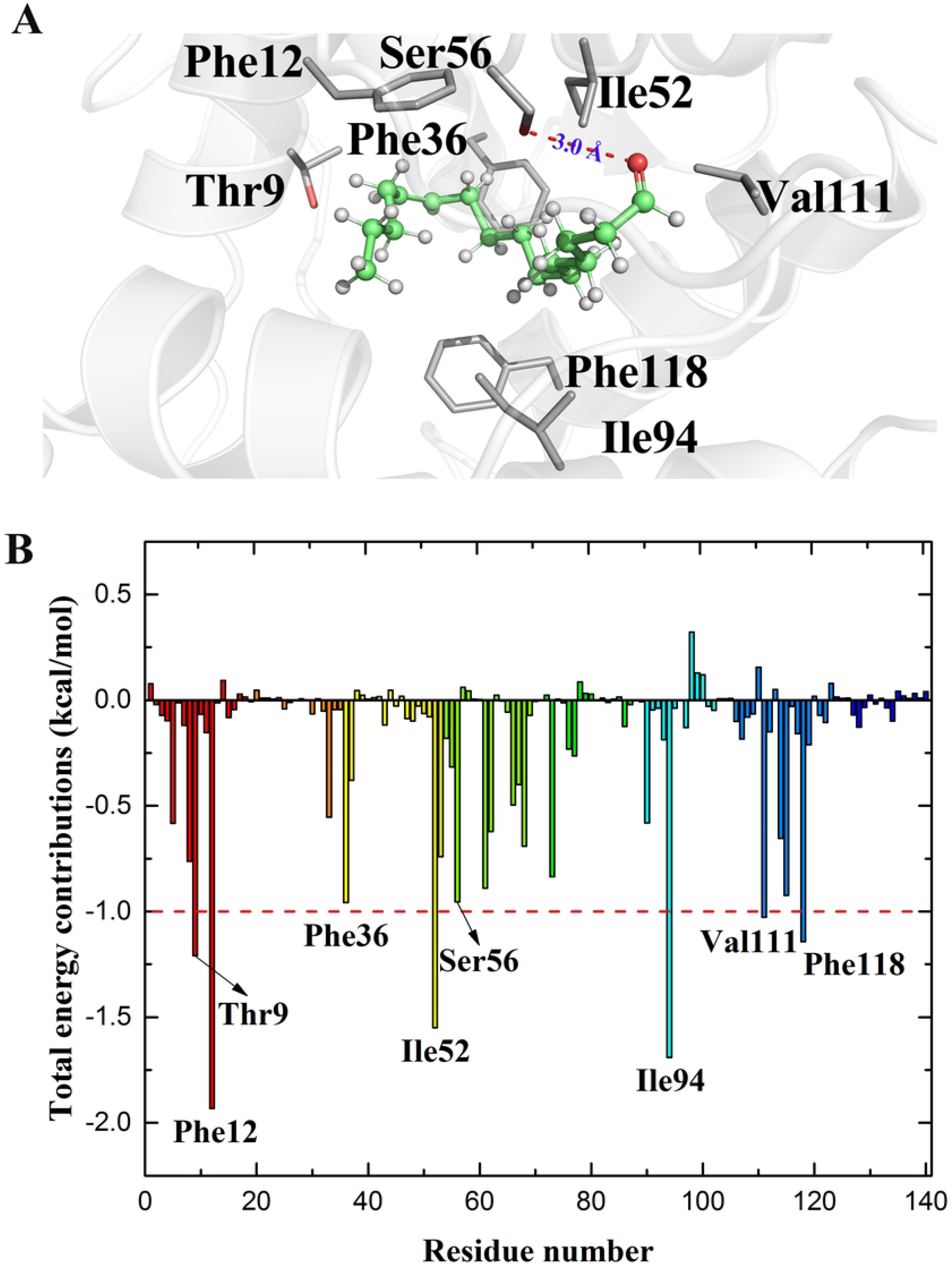
Key interactions (A) and energy contributions (B) of residues at the binding sites of the MsGOBP2-*Z*11-16: Ald complex. In (A), residues are shown as the stick model, *Z*11-16: Ald is shown as the stick-sphere model. Color code: red sphere, oxygen atom; green sphere, carbon atom; gray sphere, hydrogen atom; red dashed line, hydrogen bond. In (B), the Y-axis denotes the total binding free energy contributions of each residue in the MsGOBP2--*Z*11-16: Ald complex, the X-axis represent the residues of MsGOBP2. Residues composing the binding pocket of MsGOBP2 are marked.

Binding free energy was generally composed of three items including ΔE_VDW_, ΔE_ELE_ and solvation free energy (ΔE_SOL_). In view of this, ΔG_bind-cal_ of the MsGOBP2-*Z*11-16: Ald complex was estimated to be −5.93 ± 0.38 kcal/mol (Table 2). The yielded ΔG_bind - cal_ was fairly close (difference within −2.00 kcal/mol) to the experimental binding free energy (ΔG_bind - exp=_-7.66 kcal/mol) calculated based on the *K*_d_ between MsGOBP2 and *Z*11-16: Ald. As another dominant contributor of ΔG_bind-cal_, the ΔE_SOL_ derived from nonpolarity (ΔE_ECAVITY=_-34.15 ± 0.07 kcal/mol) was favorable to the MsGOBP2-*Z*11-16: Ald interaction (Table 1). While the ΔE_SOL_ contributed by static electricity was unfavorable, the electrostatic ΔE_SOL_ calculated by the Poisson-Boltzmann function (ΔE_EPB_) was 22.78 ± 0.26 kcal/mol (Table 1). Removing such unfavorable interactions would help to improve the affinity between MsGOBP2 and *Z*11-16: Ald. Take Ser56 for example, its sidechain E_EPB_ (S_EPB_) reached 0.76 kcal/mol (Table 2). Thus, when optimizing *Z*11-16: Ald, groups forming polar solvation interactions with Ser56 sidechain should be avoided. That coupled with the weak total E_VDW_ (T_VDW_) contributions of the two residues (Val111 and Ser56) close to *Z*11-16: Ald carbonyl drove us to suppose that introducing a methyl or other hydrophobic group to the carbon atom (C) of *Z*11-16: Ald carbonyl would optimize the MsGOBP2-*Z*11-16: Ald affinity.

## 3 DISCUSSION

It is well known that insect sex pheromones are released by female adults and accurately recognized by male adults of the same species^[40]^. Due to sexual immaturity, larvae are thought to be incapable of perceiving the sex pheromones. However, our results showed that the first two instars of *M. separata* larvae can be strongly attracted by the sex pheromone component *Z*11-16: Ald in a food context. Similar pheromone has also been reported in other lepidopteran larvae including *S. litura, S. littoralis* and *P. xylostella*^[28-30]^. These insects all responded strongly to sex pheromones of their own species in the early larval stage. We can speculate that sex pheromones left by female when laying eggs may act as an indicator of better food for the early-instar larvae^[28]^. Actually, the same semiochemical conveying different message is not unusual^[28, 41-43]^. Take (*E, E*)-8, 10-dodecadienol for example, it is not only the sex pheromone of *Cydia pomonella*^[44]^, but also acts as a synergist of *Grapholita molesta* sex pheromone^[45-47]^. For early-instar larvae, newly-hatched ones in particular, are fragile to environmental changes. It has been found that insects (e.g. *S. exempta, S. littoralis*, and *M. separata*) living in high density are more resistant to pathogens and other risks^[48-51]^, the phenomenon is termed density-dependent prophylaxis^[52]^. Thus, attracting and concentrating early-instar larvae can be taken as a self-protecting mechanism of insects.

We noticed that the positive taxis of larvae to sex pheromones became disappeared when they reached specific instars. In our tests, *M. separata* larvae exceeding the 3rd instar showed no evident response to *Z*11-16: Ald. There could be more than one reasons. On one side, the food intake of *M. separata* larvae became largely increased after entering the 3rd instar. In this way, directing them to the pheromone-containing sites would exacerbate their competition for food. On the other side, *M. separata* larvae are provided with a habit of cannibalism after the 3rd instar, the loss of taxis to *Z*11-16: Ald could avoid cannibalism effectively. When considering the cannibalism, the phenomenon of using sex pheromones as a clue to better food might not be active to cannibalistic insect species like *Helicoverpa armigera, S. frugiperda* and other related species. For *P. xylostella* larvae, their responses to sex pheromones also lost when the larvae reached the 4th instar, although no cannibalism has been observed among them^[28]^. This can be attributed to the fact that *P. xylostella* larvae at this stage are preparing for pupation and have low demand for food, after all sex pheromones are suggested as attractants for larvae only in the presence of food^[28, 29]^.

The recognition of sex pheromones is a quite complicated process where more than one proteins are involved^[1, 3, 11]^. Sex pheromones are known to be solubilized, selected and transported to the membrane-bound ORs by PBPs^[53, 54]^. Some researches using *Xenopus*-based models suggested that PBPs were not necessary, since pheromone receptors expressed in *Xenopus* oocytes can be activated by high-dosed sex pheromones in the absence of specific PBPs^[12, 25, 26, 39]^. In the larval antennae of *M. separata*, no PBPs were detected. Our work reported a sharply attenuated response to *Z*11-16: Ald in larvae expressing low level of MsGOBP2. The inefficiency of high-dosed *Z*11-16: Ald in improving the decreased response further suggested indispensability of MsGOBP2 in perceiving *Z*11-16: Ald by early-instar *M. separata* larvae. As a membrane-bound olfactory protein highly expressed in larval antennae, the MsOR3 were suggested to be in charge of *Z*11-16: Ald perception as well. The silence of *MsOR3* gene caused a significant decrease of larvae attracted by *Z*11-16: Ald. MsOR3 has been reported as a receptor specifically and abundantly expressed in the antennae of *M. separata* male adults, oocytes expressing MsOR3 was detected to give dose dependent responses to *Z*11-16: Ald^[25, 26]^. In addition to MsOR3, MsOR1 was defined as a second receptor of *Z*11-16: Ald in male adults^[39]^. However, we did not detect the expression of MsOR1 in larval antennae, indicating that MsOR1 was not involved in the perception of *Z*11-16: Ald in the early-instar larvae of *M. separata*. Combining the indispensability of MsGOBP2, it can be concluded that the recognition of *Z*11-16: Ald in *M. separata* larvae was mediated by the interplay between MsGOBP2 and MsOR3. When considering the slight reduction of *Z*11-16: Ald-based food preference in *MsGOBP1*-silenced larvae, it is possible that MsGOBP1 act as an auxiliary protein in *Z*11-16: Ald perception. As for MsOR2, it has been reported to show dose dependent responses to another (*Z*)-9-tetradecenal (*Z*9-14: Ald), a second component of *M. separata* sex pheromone^[25, 38]^. While in our research, the silence of MsOR2 increased the number of larvae that made no choice between maize leaves with and without *Z*11-16: Ald, suggesting that MsOR2 may play a role in the feeding of *M. separata* larvae. Based on these discoveries, a schematic diagram describing the role of MsGOBPs and MsOR3 were provided (Figure 6). It is shown that the hydrophobic molecule *Z*11-16: Ald was captured and ferried by the soluble MsGOBP2 to activate MsOR3 directly or indirectly.

**Figure 6.**
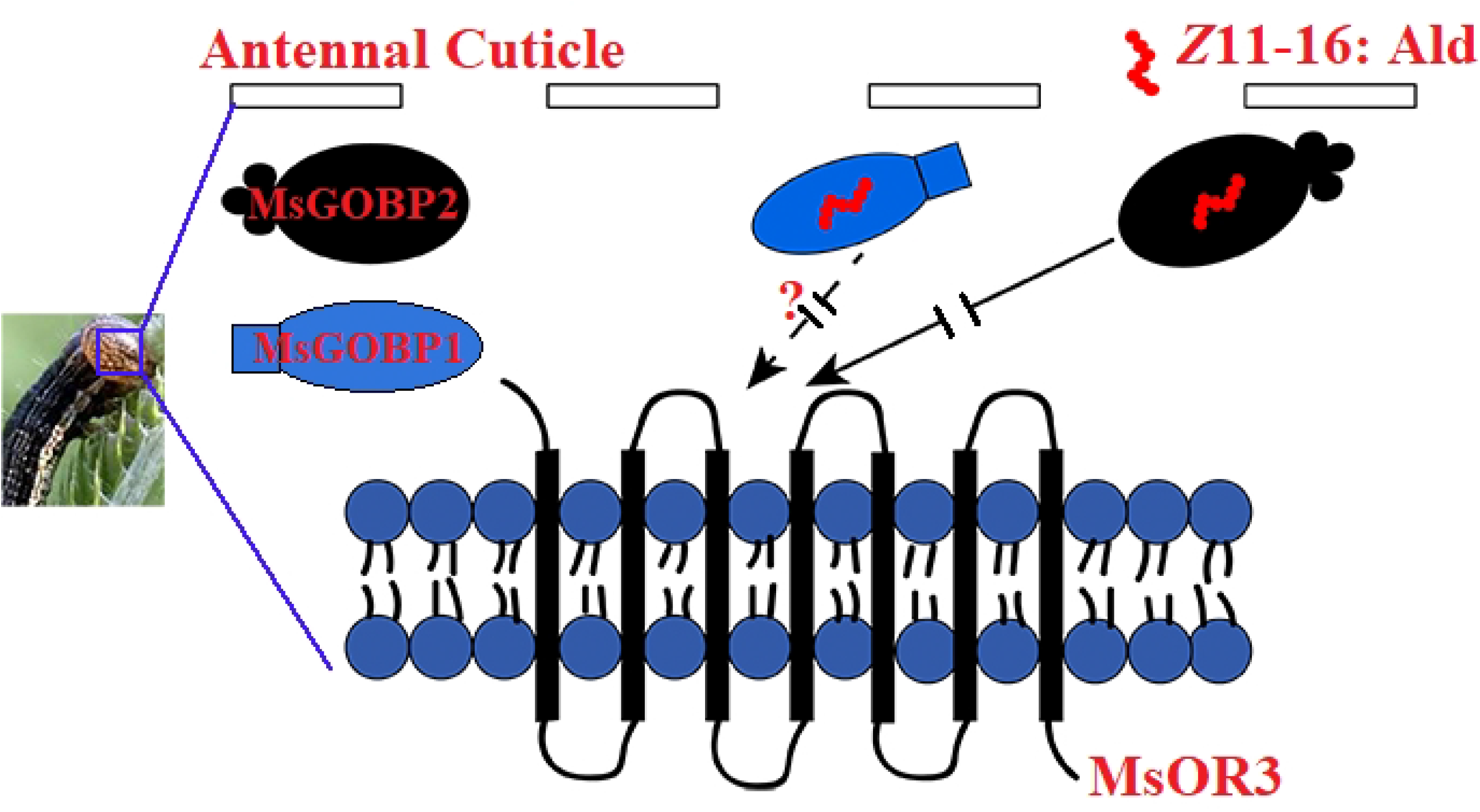
The schematic diagram of *Z*11-16: Ald perception i*n Mythimna separata* larvae.

Relative to the adult system for *Z*11-16: Ald perception, the same system for larvae was quite simple, involving only MsGOBP2 and MsOR3 (Figure 6). In light of the relatively simple chemoreception system, lepidopteran larvae showing responses to their sex pheromones can be developed as suitable models for the *in vivo* exploration of sex pheromone perception.

In summary, we examined the responses of *M. separata* larvae to the sex pheromone component *Z*11-16: Ald and found that the early-instar larvae exhibited evident preference for the host spiked with *Z*11-16: Ald. Combined RNAi and larval behavior tests revealed that MsGOBP2 and MsOR3 were two olfactory proteins in charge of *Z*11-16: Ald perception in the early-instar larvae of *M. separata*. A *Z*11-16: Ald perception mechanism mediated by the interplay between MsGOBP2 and MsOR3 was consequently proposed. As the first soluble protein interacting with *Z*11-16: Ald in larval antennae, MsGOBP2 showed high affinity to *Z*11-16: Ald. The interactions between MsGOBP2 and *Z*11-16: Ald was also depicted. The present work demonstrated the influences of *Z*11-16: Ald on the host-feeding choice regulation of *M. separata* larvae, and elucidated a putative mechanism for the perception of *Z*11-16: Ald in *M. separata* larvae. Related discoveries not only suggested the applicability of lepidopteran larvae as *in vivo* models to explore mechanisms for sex pheromone perception, but also provided basis for developing olfaction-based management of *M. separata* larvae which is the most destructive stage throughout their whole life history.

## 4 MATERIALS AND METHODS

### 4.1 Insects and chemicals

The *M. separata* population was provided by the Key Lab of Plant Protection Resources & Pest Management of Ministry of Education, Northwest A&F University. Larvae were reared with fresh maize leaves in the condition of 23±1 °C, RH 60%, and a photoperiod of 16L: 8D. The pheromone (*Z*)-11-hexadecenal (*Z*11-16: Ald), the fluorescence probe N-phenyl-1-naphthylamine (1-NPN) and the GC-grade methanol were purchased from Aladdin (Shanghai, China).

### 4.2 Larval behavior tests

To test the influence of *Z*11-16: Ald on the food selection of *M. separata* larvae, a diet choice assay was conducted using a petri dish (diameter=15 cm). In the petri dish, two pieces of fresh maize leaves (1.5□×1.5 cm), one added with 10 ng *Z*11-16: Ald in methanol and the other with equivalent amount of methanol, were placed diametrically opposite (Figure 1A). The 1st-4th instar larvae of *M. separata* were treated with starvation for 8 h. For each instar, ten replicates were performed, ten larvae were used for each replicate. When the test begins, ten starved larvae were positioned at the release point, the larvae numbers on or close to each leaf piece were counted after 10 min. The attractivity of *Z*11-16: Ald alone to *M. separata* larvae was tested in the same way, with fresh maize leaves being removed.

Then, the diet choice assay was conducted on maize seedlings using the 2nd instar larvae. As shown in Figure 1C, two maize seedlings (two euphylla) were planted in a pot (diameter=20 cm), with a seedling distance of 10 cm. One seedling was sprayed with 100 ng *Z*11-16: Ald in methanol, the other was sprayed with equivalent amount of methanol. Twenty starved larvae collected in a petri dish were placed in the midpoint between the two seedlings, the larvae numbers on o each seedling were counted after 30 min. Ten replicates were performed for the test.

We also tested the attractivity of non-host plants sprayed with the mixture of *Z*11-16: Ald and crude extracts of maize leaves. Pea seedlings and maize seedlings were planted in a pot (diameter=20 cm), with a seedling distance of 10 cm. the pea seedlings were sprayed with the mixture of *Z*11-16: Ald and crude extracts of maize leaves. Twenty starved larvae collected in a petri dish were placed in the midpoint between the two seedlings, the larvae numbers on o each seedling were counted after 30 min. Ten replicates were performed for the test.

All experiments were conducted between 09:00 and 16:00 under the same laboratory condition (uniform light and constant temperature at 25 °C).

### 4.3 Expression profiles of several olfactory proteins

The reverse-transcription PCR (RT-PCR) was adopted to test the expression profiles of 10 *M. separata* olfactory genes including *MsPBP* (GenBank: AB263112), *MsPBP1* (GenBank: MH168089), *MsPBP2* (GenBank: MH168090), *MsGOBP1* (GenBank: MH175135), *MsGOBP2* (GenBank: MH175137), *MsPR1* (GenBank: MK500698), *MsPR2* (GenBank: MK500699), *MsOR1* (GenBank: AB263110), *MsOR2* (GenBank: MH717242) and *MsOR3* (GenBank: MH717241) in larvae. Total RNA was extracted from the head of the 1st-4th instar larvae of *M. separata* and subjected to cDNA synthesis. Integrity of cDNA templates was tested using the *M. separata β-actin* gene (*Msβ-actin*, GenBank: GQ856238) as the reference gene. Specific primers (Table S1) for the 10 genes were designed according to their sequences downloaded from GenBank. The RT-PCR was conducted under the following conditions: 95 °C for 3 min, 35 cycles of 95 °C for 30 s, 58 °C for 30 s, and 72 °C for 1 min, then a final incubation at 72 °C for 5 min. The PCR products were purified and sequenced for genes confirmation.

### 4.4 Protein expression and purification

The *MsGOBP1* and *MsGOBP2* genes were sub-cloned into the pColdII vector (TaKaRa, Japan) and successfully transformed into the competent TransB strains (Transgen, China). The yielded strains were inoculated in 1000 mL LB medium (Amp+) and cultured till the logarithmic phase (OD_600_=0.6) under the condition of 37 °C, 200 rpm. The cultures were then precooled to 15 °C, added with 0.6 mM Isopropyl-β-D-Thiogalactoside (IPTG), and further incubated for 20 h (15 °C, 160 rpm) to induce the expression of MsGOBP1 and MsGOBP2 proteins. Thereafter, the strains were collected by centrifugation (5000 *g*, 4 °C, 10 min) and incubated with lysozyme for 30 min at room temperature before being disrupted by ultrasonication. The yielded periplasmic fractions were subjected to centrifugation (12000 *g*, 4 °C, 30 min) to separate the sediments completely from the supernatants. The supernatants were purified by the Ni^2+^-NTA column (Qiagen, Germany) and dialyzed against 10 mM PBS buffer (pH 7.4). After that, the purified MsGOBP1 and MsGOBP2 proteins were digested by Enterokinase (Solarbio, China) to remove the expression tag, re-purified by the Ni^2+^-NTA column (Qiagen, Germany), and finally dialyzed against 10 mM PBS buffer (pH 7.4). Detailed procedures for the expression and purification of the two proteins were similar to that of other OBPs in our former reports^[55-57]^. The produced MsGOBP1 and MsGOBP2 proteins can be directly used for prospective *in vitro* binding assay.

### 4.5 *In vitro* binding assay

Fluorescence competitive binding assay is a common method to evaluate the *in vitro* binding affinity between ligands and insect OBPs. We initially titrated the 2 μM MsGOBP1/MsGOBP2 solutions with the fluorescent probe, N-phenyl-1-naphthylamine (1-NPN), yielding the dissociation constants between the two proteins and 1-NPN (*K*_1-NPN_). To measure the in vitro binding affinity between *Z*11-16: Ald and MsGOBP1/MsGOBP2, solutions containing 2 μM MsGOBP1/MsGOBP2 proteins and 2 μM 1-NPN were titrated with 1 mM *Z*11-16: Ald. Each trial was performed in three repetitions. All fluorescence responses were detected on a Hitachi F-2700 spectrofluorimeter (Hatachi, Japan), with excitation wavelength being set as 337 nm and emission spectra being recorded between 370 nm and 500 nm. The dissociation constants between MsGOBP1/MsGOBP2 and *Z*11-16: Ald (*K*_d_) were calculated based on the fluorescence intensity changes and the values of *K*_1-NPN_^[58]^.

### 4.6 RNAi of genes encoding olfactory proteins

The primer pairs for double-stranded RNA (dsRNA) were designed based on the cDNA sequences of genes encoding five olfactory proteins including MsGOBP1, MsGOBP2, MsPR1, MsOR2 and MsOR3 (Table S1). Each dsRNA of the *MsGOBP1, MsGOBP2, MsPR1, MsOR2* and *MsOR3* genes (*dsGOPB1, dsGOBP2, dsPR1, dsOR2* and *dsOR3*) was synthesized using the T7 RiboMAX Express RNAi system (Promega, America). The newly-hatched *M. separata* larvae were continuously fed with artificial diet containing dsRNA (1 μg dsRNA/1 g artificial diet) for 72 h to suppress the expression of *MsGOBP1, MsGOBP2, MsPR1* and *MsOR3* genes in larval antennae (59-61). The interference efficiency of each gene was evaluated by real time qPCR (RT-qPCR). The treated larvae were then subjected to behavioral tests which were performed as mentioned above. For RNAi, the RNase-free water and dsRNA of GFP (*dsGFP*) were fed as control. For RT-qPCR, the *Msβ-actin* was used as reference gene. All experiments were repeated three times.

### 4.7 Structural evidences for the high affinity between Z11-16: Ald and MsGOBP2

The 3D structure of the MsGOBP2 protein was produced in the Modeller9.10 software^[62]^. The molecule *Z*11-16: Ald was docked into the binding cavity of MsGOBP2 using the GOLD2020.3.0 software^[63]^. The yielded MsGOBP2-*Z*11-16: Ald complex was subjected to Amber12 package for molecular dynamics (MD) simulations^[64]^. It is based on the representative conformation produced by MD simulations that the structural evidences for the high affinity between MsGOBP2 and *Z*11-16: Ald were discussed. Binding free energy calculation of the MsGOBP2-*Z*11-16: Ald complex was calculated by the Molecular Mechanics-Poisson-Boltzmann surface area (MM-PBSA) method in the Amber12 package^[65-67]^. Per-residue free energy decomposition of the MsGOBP2-*Z*11-16: Ald system was performed using the Molecular Mechanics-Generalized-Born surface area (MM-GBSA) method in Amber12 package^[68]^. Detail processes were provided in the Supporting Information (SI).

### 4.8 Statistical analysis

All data were analyzed by the Graphpad prism 6 software. A χ^2^ test was used to compare the larvae number of final choices made between the treatment and the control. The relative expression levels of *MsGOBP1, MsGOBP2, MsPR1, MsOR2* and *MsOR3* were calculated based on the 2^-ΔΔCt^ method, and analyzed using one-way ANOVA followed by Tukey’s multiple comparison test.

The *K*_d_ value between MsGOBP2 and *Z*11-16: Ald was calculated using equation (1):

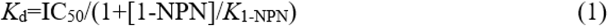

In equation (1), IC_50_ is the concentration of *Z*11-16: Ald quenching half of the fluorescence intensity; [1-NPN] is the concentration of free 1-NPN; *K*_1-NPN_ is the dissociation constant between MsGOBP2 and 1-NPN.

## ACKNOWLEDGEMENTS

This work was supported by the Chinese Universities Scientific Fund (Z1010422004), the National Natural Science Foundation of China (31801797), the Chinese Universities Scientific Fund (2452018008), the Natural Science Foundation of Jiangsu Province (BK20180902) the Sci-Tech Planning Project of Yangling Demonstration Zone (2018NY-02), the National Natural Science Foundation of China (21503272).

## AUTHOR CONTRIBUTIONS

Z.T. and J.Y.L. conceived the project. Z.T. and J.Y.L. designed the experiment. J.Y.L., Z.T., S.C.C., T.Z., and R.C.L. performed the experiments and prepared the manuscript. Z.T. supervised the study and contributed the reagents and materials. All authors contributed to data analysis.

## SUPPORTING INFORMATION

Supplements to the methods of homology modeling, molecular docking, molecular dynamics simulations, binding free energy calculation, per-residue free energy decomposition; Supplements to results; Supporting figures (Figure S1: Z11-16: Ald alone showed no attractivity to the 1st-3rd instar larvae of *Mythimna separata*; Figure S2: The host-feeding choice of the 2nd instar *Mythimna separata* larvae between maize seedlings and pea seedlings sprayed with crude extracts of maize leaves; Figure S3: SDS-PAGE analysis of the two recombinant MsGOBPs; Figure S4: Dose-response curves for 1-NPN against the two recombinant MsGOBPs; Figure S5: The structure of MsGOBP2; Figure S6: Ramanchandran plot of the generated MsGOBP2 model; Figure S7: 3D/1D profile of the generated MsGOBP2 model; Figure S8: Molecular dynamics (MD) simulations of the MsGOBP2-*Z*11-16: Ald complex; Figure S9: Cluster analysis of the MD trajectories produced during the 100 ns MD simulations of the MsGOBP2-*Z*11-16: Ald complex); Supporting tables (Table S1: Primers used in the experiment; Table S2: Cluster analysis of MsGOBP1-Z11-16: Ald complexes based on the MD simulations trajectories)

